# Refined mechanism of *Mycoplasma mobile* gliding based on structure, ATPase activity, and sialic acid binding of machinery

**DOI:** 10.1101/823039

**Authors:** Miyuki S Nishikawa, Daisuke Nakane, Takuma Toyonaga, Akihiro Kawamoto, Takayuki Kato, Keiichi Namba, Makoto Miyata

## Abstract

*Mycoplasma mobile*, a fish pathogen, glides on solid surfaces by repeated catch, pull, and release of sialylated oligosaccharides by a unique mechanism based on ATP energy. The gliding machinery is composed of huge surface proteins and an internal jellyfish -like structure. Here, we elucidated the detailed three-dimensional structures of the machinery by electron cryotomography. The internal tentacle -like structure hydrolyzed ATP, which was consistent with the fact that the paralogs of the α- and β-subunits of F_1_-ATPase are at the tentacle structure. The electron microscopy suggested conformational changes of the tentacle structure depending on the presence of ATP analogs. The gliding machinery was isolated and shown that the binding activity to sialylated oligosaccharide was higher in the presence of ADP than in the presence of ATP. Based on these results, we proposed a model to explain the mechanism of *M. mobile* gliding.

**IMPORTANCE:** The genus *Mycoplasma* is made up of the smallest parasitic and sometimes commensal bacteria; *Mycoplasma pneumoniae*, which causes human walking pneumonia, is representative. More than ten *Mycoplasma* species glide on host tissues by novel mechanisms always in the direction of the distal side of the machinery. *Mycoplasma mobile*, the fastest species in the genus, catches, pulls, and releases sialylated oligosaccharides (SOs), the carbohydrate molecules also targeted by influenza viruses, by means of a specific receptor and using ATP hydrolysis for energy. Here, the architecture of the gliding machinery was visualized three-dimensionally by electron cryotomography (ECT), and changes in the structure and binding activity coupled to ATP hydrolysis were discovered. Based on the results, a refined mechanism was proposed for this unique motility.

## (INTRODUCTION)

Mycoplasmas are parasitic and occasionally commensal bacteria that lack peptidoglycan layers and have small genomes (1, 2). *Mycoplasma mobile*, a fish pathogen, has a membrane protrusion, a gliding machinery at one pole and glides in the direction of the protrusion (Fig. 1A) (3-6). The average speed is 2.0 to 4.5 μm/s, or 3 to 7 times the cell length /s, with a propulsive force up to 113 pN (Movie S1) (7-10). This motility, combined with the ability to adhere to the host cell surface, likely plays a role in infection, as has been suggested for another species, *Mycoplasma pneumoniae* (4, 11, 12). The motor proteins involved in this motility are unlike the motor proteins involved in any other form of bacterial or eukaryotic cell motility (3-6, 13, 14).

**FIG 1.**
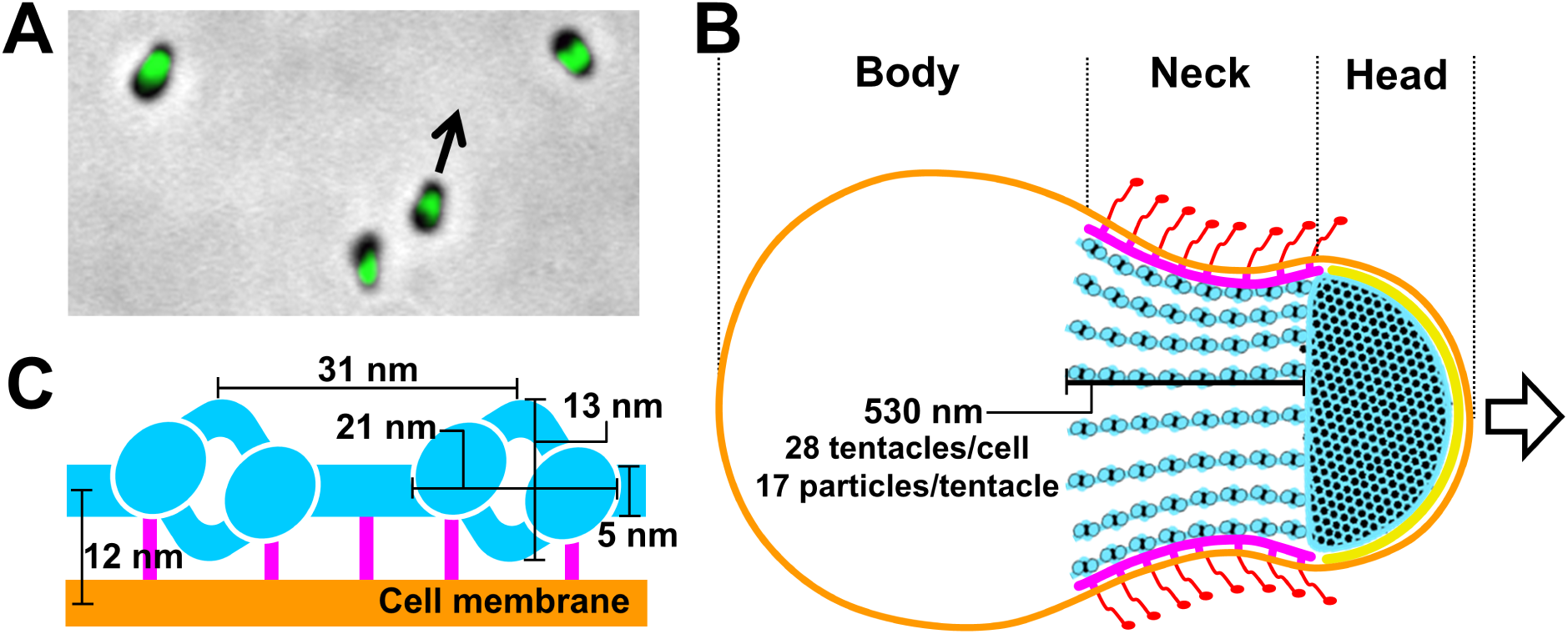
Cell and the gliding machinery of *M. mobile*. (A) Optical microscopy of cells. Phase-contrast image was overlaid by green fluorescence image labeled for MMOB1660, a protein involved in the gliding machinery. The cells are always gliding in the direction indicated by a black arrow. A related movie is available (Movie S1). (B) Schematic illustration of whole cell architecture. The cell can be divided into head, neck, and body from the front. The cell membrane (orange), jellyfish-like structure (bell + tentacles)(blue), undercoats at head region (yellow), undercoats at neck region (pink), and legs (red) are presented using the common colors with Fig. 2G, based on the results presented in Fig. 2 and a previous study (3). Twenty eight tentacles are present in a cell. They are 530 nm long and contain 17 particles. (C) Magnified schematic of tentacle part of jellyfish-like structure with cell membrane and dimensions, based on the results presented in Fig 2. Jellyfish-like structure (blue), bridges at the neck region (pink), and cell membranes (orange) are presented using the common color. Note that the only bridge part is colored pink.

The cell surface can be divided into three parts beginning at the front end, i.e., the head, neck, and body (Fig. 1B) (3, 5, 6, 15-17) Three large proteins, Gli123, Gli349, and Gli521, with respective masses of 123, 349, and 521 kDa, are involved in this gliding mechanism and are localized exclusively at the cell neck (7, 15, 16, 18-21). About fifty-nanometer legs corresponding to Gli349 can be seen protruding from the neck surface by electron microscopy (EM) (22, 23). The internal structure of the machinery is a “jellyfish” - like structure composed of 10 jellyfish-structure proteins (JSPs) (22, 24, 25). Interestingly, the amino acid sequences of MMOB1660 and MMOB1670, belonging to JPSs show high similarity to the respective α- and β-subunits of F_1_-ATPase, the catalytic subunit of proton pumps. However, these proteins are unlikely to function in an F-type proton pump because F-type proton pumps require additional subunits (26) and the *M. mobile* genome has another locus containing a complete set of pump subunits. The energy for motility should be supplied through ATP, although gliding can be driven also by GTP and dATP (27-30), and the direct binding targets for gliding are the sialylated oligosaccharides (SOs) found on the surface of animal tissue (31-36). On the basis of the above information, we proposed a working model called the centipede or power stroke model, in which the cells are propelled by a foot, a part of Gli349 that, through repeated cycles driven by the hydrolysis of ATP, catches and releases SOs (3, 5, 7, 37). However, the force generation by ATP hydrolysis in the cell body and the subsequent transmission to the cell surface are still unclear. Elucidation of the detailed three-dimensional composition of the gliding machinery will be crucial in order to fully clarify the gliding mechanism.

In this study, we analyzed the structure of the gliding machinery by electron cryotomography (ECT) and studied the reactions involved in ATP hydrolysis; based on our findings, we proposed a mechanism for *M. mobile* gliding.

## (RESULTS

### (Detailed structures of the gliding machinery observed by ECT

To visualize the gliding machinery three-dimensionally, intact cells (Fig. 2A, Movie S2), permeabilized cells (Fig 2B and C, Movie S3), and the internal jellyfish-like structures (Fig. 2D, Movie S4) were analyzed by ECT. Cultured cells were collected, suspended in a growth medium, put on a grid, blotted with paper filters, frozen quickly, and observed by ECT. The contrast of the internal structures in an intact cell was low because the thick and dense cell bodies inhibited transmission of the electron beams (Fig. 2A). To solve this problem, we prepared permeabilized cells using 0.7% Tween 20 (Fig. 2B and C), and then succeeded in observing filamentous structures on the cell surface, with a length ranging from 60 to 100 nm and featured with a thick end. (Fig. 2E left and Fig. S1A). Two types of filamentous structures were also observed as undercoats beneath the cell membranes of the head and neck regions (Fig. 2C middle, E middle and right). Moreover, a novel structure was found to bridge the filament to the cell membrane for a distance of 12.0 ± 1.3 nm (*n* = 144 positions in 16 cells) (Fig. 2E right and Fig. S1B(i)). Although the jellyfish-like structure could be observed in the permeabilized cells, the contrast of the jellyfish-like structure was not sufficient for further analyses (Fig. 2C). Therefore, we removed the cell membrane by treatment with 0.08% Triton X-100, resulting in a clearer image of the jellyfish-like structure (Fig. 2D and F). On the basis of these observations, we concluded that each cell has 28 ± 3 tentacle structures (*n* = 15 cells)(Fig. S1B(ii)), each 530 ± 110 nm long (*n* = 87 tentacle structures)(Fig. S1B(iii)), and that the tentacle structures contain globular particles with a clear periodicity of 32 ± 2 nm (*n* = 13 tentacle structures). A three dimensionally rendered image of permeabilized cell is presented in Fig. 2G and Movie S5. The undercoats and the tentacle structures were analyzed for periodicity by image averaging and its Fourier transform (Fig. 2H). Common periodicities were observed in the Fourier transforms of the neck undercoat and the tentacle structure but not in the front undercoat, suggesting that the first two are derived from a common structure. The tentacle image was obtained as an average of 199 images of 23.6 nm thick slice, featured with 31-nm repeating units including a particle 13 nm wide and 21 nm long connected by axial stalks 5 nm in diameter and featured with lateral protrusions at two positions (Fig. 2H). The neck undercoat image was obtained as an average of 61 images featured with possible projections to the cell membrane. The structures and their dimensions are schematically summarized in Fig. 1B and C.

**FIG 2.**
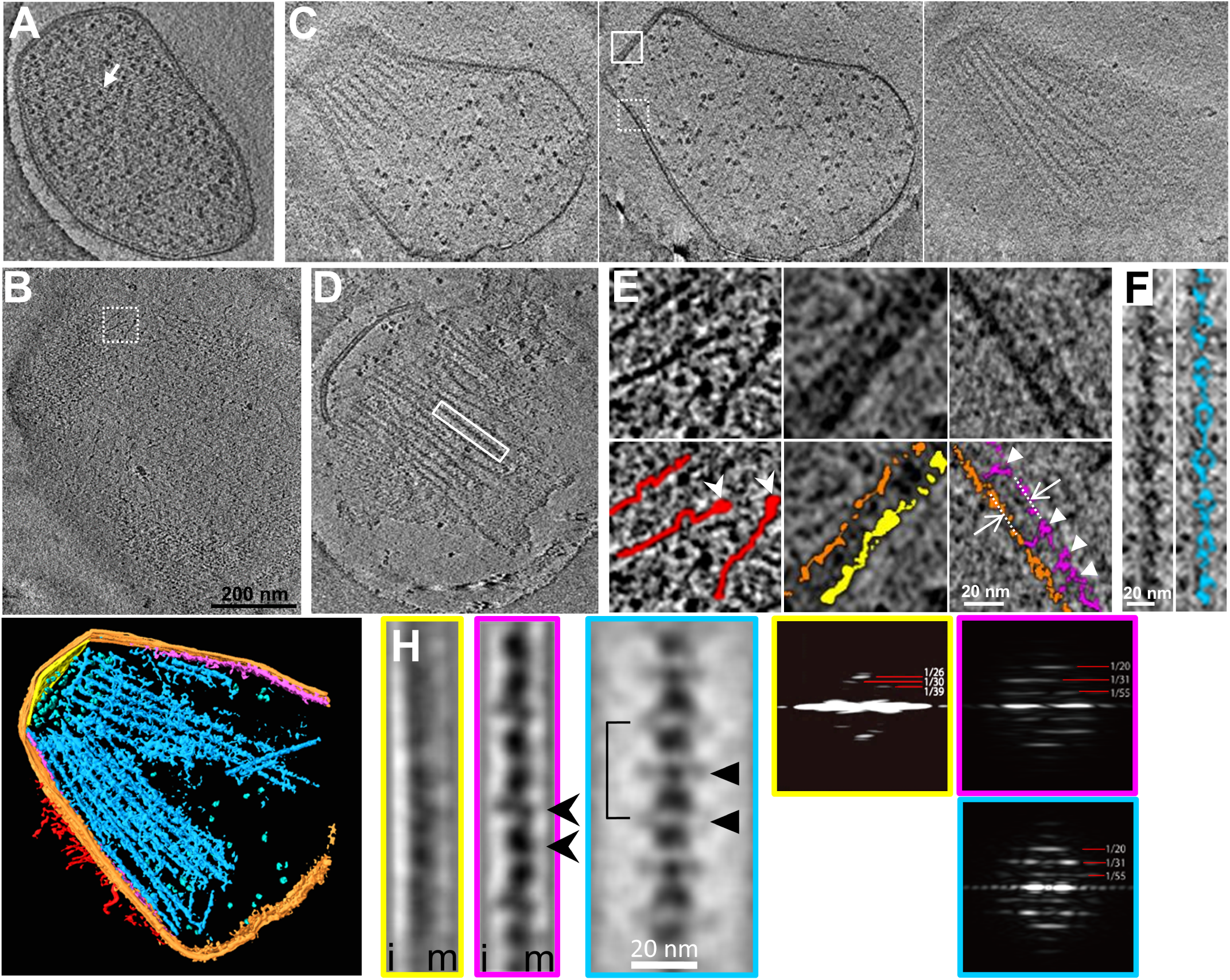
Detailed structures of the gliding machinery observed by ECT. (A) Slice image of an intact cell including many ribosomes marked by a white arrow. (B) Slice image at the surface of a permeabilized cell. (C) Three slice images of a permeabilized cell at different heights. (D) Slice image of the jellyfish-like structure. Slice images (A) to (D) are also available as Movies S2 to S5, respectively. The same magnification was applied to (A) to (D). (E) Magnified images of the slices. Original and colored images are presented in the upper and lower panels, respectively. The dashed-boxed area in (B), and solid- and dashed-boxed areas in (C) are magnified in left, middle, and right columns, respectively. The surface filamentous structures, cell membrane, and undercoats at the head and neck membranes are colored red, orange, yellow, and purple, respectively. The thick end of filamentous structure on surface is marked by a white arrow head. The distance between tentacle structures and the cell membrane is shown by white arrows. The connecting points are marked by white triangles. (F) Magnified image of the solid-boxed area in (D). The tentacle structure is colored blue in the right panel. The slice images (A) to (F) are 2.36 nm thick. (G) Three dimensional image rendered for 146 nm-thick slice of a tomogram of a permeabilized cell reconstructed by ECT. The ribosome is colored green. Other colors are common in Fig 2. (H) Averaged images (left) and their Fourier transforms (right) of periodical structures. The images of the undercoat at the head (yellow), undercoat at the neck (pink), and tentacle structures (blue) were integrated from 32, 61, and 199 images, respectively. The slice image is 23.6 nm thick. Membrane and inner sides relative to the undercoats are marked (m) and (i), respectively. Bridges, repeat unit and lateral protrusions are marked by a black arrow head, a solid line, and a black triangle, respectively. Strong layer lines in the Fourier transforms are marked with their spacings in 1/nm.

### The jellyfish-like structure hydrolyzes ATP with conformational changes

As the jellyfish-like structure contains the paralogs of F_1_-ATPase catalytic subunits, we assayed the ATPase activity of the isolated jellyfish-like structure at room temperature (RT), and found that the structure hydrolyzes ATP with *K*_*m*_ of 66 µM (Fig. 3A). If this activity is derived from the β-subunit paralog, MMOB1670, the maximum turnover rate is calculated to be 0.09 molecule /s, based on the estimation of protein amount from SDS-PAGE analysis. Considering that the amino acid sequences suggested that the other protein components of the jellyfish-like structure do not exhibit ATPase activity (22), MMOB1670 should hydrolyze ATP. This ATPase activity was inhibited by addition of sodium azide. The *K*_*m*_ and the maximum turnover rate were calculated to be 84 and 76 µM, and 0.063 and 0.033 molecule /s under 15.4- and 154-mM sodium azide, respectively.

**FIG 3.**
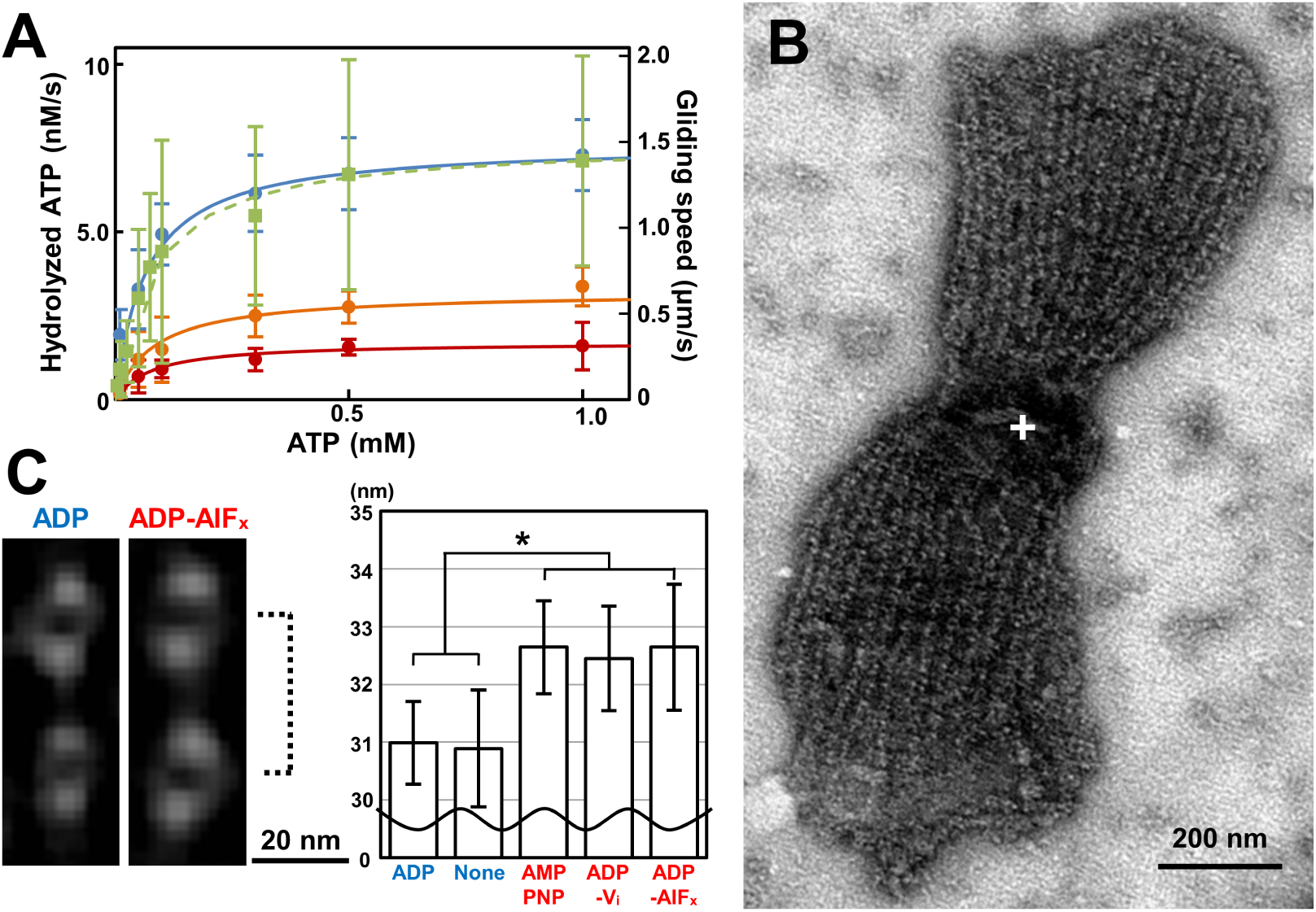
Reactions of the jellyfish-like structure with ATP. (A) ATPase activity of the isolated jellyfish-like structure and speed of the gliding heads under various ATP and sodium azide concentrations. The ATPase activities under 0-, 15.4-, and 154-mM sodium azide are indicated by blue, orange, and red filled circles respectively, and the gliding speeds without sodium azide are indicated by solid squares, and these data were fitted as solid and broken lines, respectively for ATPase activities and gliding speeds by Michaelis-Menten kinetics with *K*_m_ values of 66 μM and 60 μM, and *V*_max_ values of 7.6 nM/s and 1.4 μm/s. The concentration of ATPase β-subunit paralogs was estimated as 82 nM and the maximum turnover rate was estimated as 0.09 /s per β-subunit paralog. (B) Negatively-stained EM image of the jellyfish-like structure in the presence of ATP. The bell structure is marked by a white cross. (C) Averaged images (left) and graph of interval lengths (right) of the tentacle structures. The interval lengths of tentacle structures shown by a broken line in the presence of various ATP analogs are presented in the graph. The measurement process is shown in Fig. S2.

Next, the possibility of conformational changes in the jellyfish-like structure was examined by averaging the EM images in the presence of various ATP analogs (Fig. 3B – C, S2). Two particles on a linear tentacle structure in the presence of each of AMPPNP, ADP-V_i_, ADP-AlF_x_ and ADP, and absence of nucleotide (None) were selected, and integrated into nine classes from 108, 101, 109, 102, and 109 images, respectively. The distance between the minimum peaks of particle in density profile was measured and averaged for the nine classes in each nucleotide state. The interval of the particles was about 2 nm longer under the conditions that included AMPPNP, ADP-V_i_, and ADP-AlF_x_ compared to the interval in the presence of ADP and the absence of nucleotides. The significance of these differences was approved by Student’s *t*-test (*: *p* < 0.05). These results show that the structural change occurs at the tentacle structure, depending on the ATP hydrolyzing forms.

### Preparation of the gliding head

To examine coupling between SO binding and ATP hydrolysis, we intended to isolate the gliding machinery. In a previous study, *M. mobile* cells were elongated by treatment with detergents and their gliding was analyzed (8). In the present study, we found that, in the elongated cells, the small portion of the back end of the cell (i.e., the end opposite the gliding direction) that begins to pull away from the main body of the cell rarely fully detaches and glides away (Fig. S3A), a result which has also been observed in *M. pneumoniae*, and may arise from the same mechanism (38). Basically, an *M. mobile* cell usually has only one gliding organelle at a pole, but during the preparation for cell division, it sometimes has two gliding machineries at both poles (39, 40). The gliding force directing to both sides occasionally generates tension in the elongated cell, resulting in the detachment of one machinery from the other part. Since the pole in the gliding direction is named the head, we call the detached machinery the gliding head. The above observation showed that the small part of the cell can glide without the other parts, suggesting that it might be possible to artificially isolate the gliding machinery. To investigate this possibility, we elongated the cultured cells by treatment with 0.1% Tween 60, then sonicated and subjected them to sucrose-gradient centrifugation, and isolated fraction containing the gliding heads (Fig. 4A and Movie S6). The gliding heads were reactivated for gliding by placing them on an SOs-coated glass slide and adding ATP, as described previously for a permeabilized cell model called the gliding ghost (27-29). Immunofluorescence microscopy, Western blotting, and peptide mass fingerprinting (PMF) demonstrated that the gliding heads retained Gli521, Gli349, Gli123, MMOBs 1650, 1670, 4530, 1660, and 1620 which should be involved in the gliding machinery (Fig. 4B and Figs. S3B, S4A). The negatively-stained EM image showed that most of the cell pieces in the fraction contained the gliding machinery based on the observation of the internal jellyfish-like structure. The machinery could be observed more easily in the gliding head than in the intact cells due to the removal of the cytosol, which improved the image contrast compared to the intact cells. However, the contrast was not better than that of the permeabilized cells, because the structures of the gliding heads were still densely packed.

**FIG 4.**
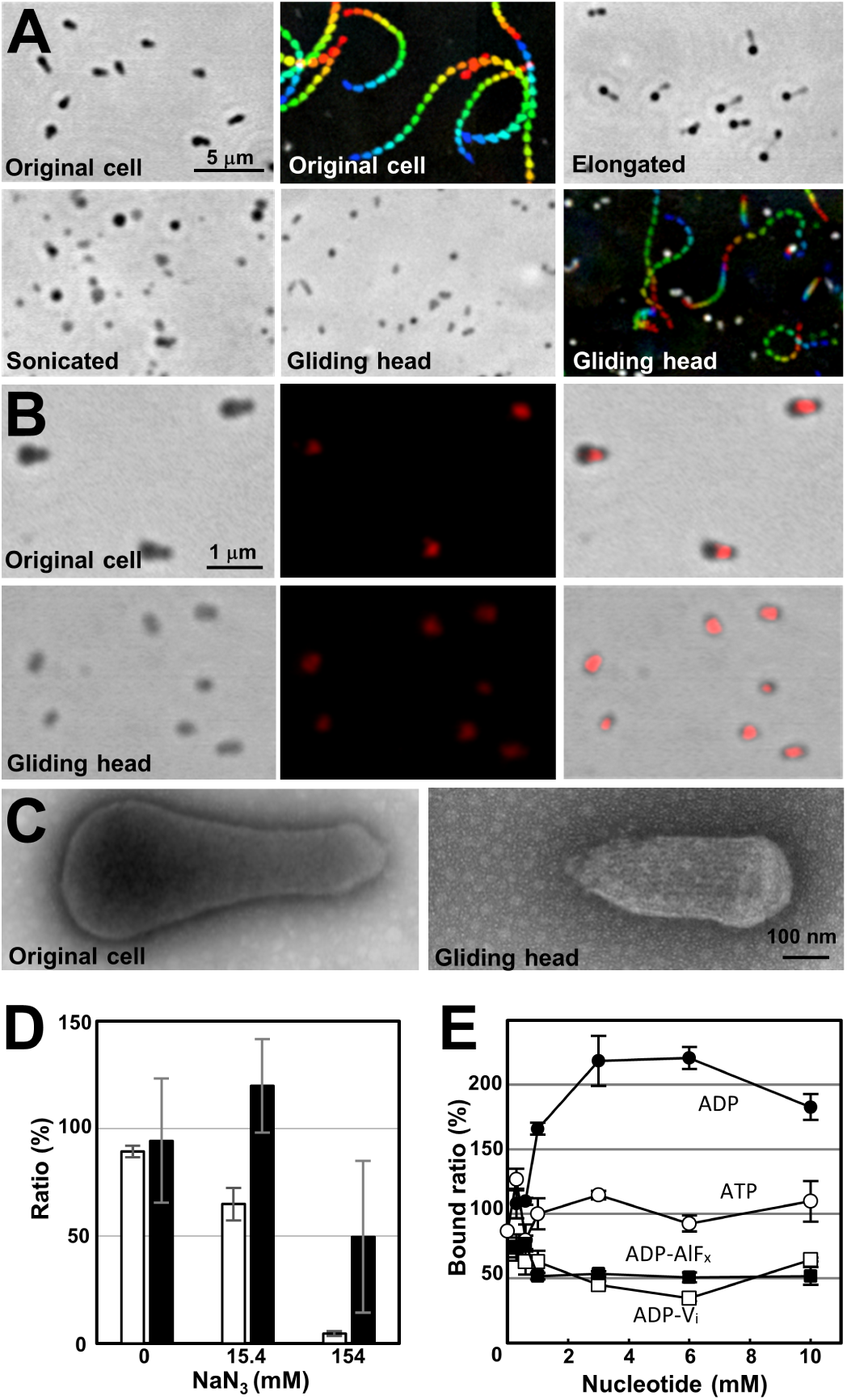
Isolation and SO binding of the gliding heads. (A) Isolation and reactivation of the gliding heads traced by phase-contrast microscopy. The original cells were elongated and sonicated, and the gliding heads were isolated. The heads were reactivated for gliding by the addition of 2 mM ATP. This gliding movie is available as Movie S6. Gliding motility is presented as an integrated movie image, where movie frames at intervals of 0.5 s are colored from purple to red, and integrated for 10 s. (B) Localization of gliding proteins examined by immunofluorescence microscopy. The images of an original cell and gliding head are presented in the upper and lower panels, respectively. Fluorescent signals labeled for Gli349 (middle) were overlaid on the phase-contrast images (left), and presented as merged images (right). (C) Negatively-stained EM images of the original cell and gliding head. (D) Inhibition on binding and gliding speed of gliding heads by sodium azide. The numbers of bound gliding heads at 50 s after addition of sodium azide were counted for three independent fields including 60-119 gliding heads and presented as ratios to the numbers before the addition by white bars with SDs. The gliding speeds were measured for 11 gliding heads and presented as ratios to those before the addition by black bars with SDs. (E) Binding of the gliding heads to a glass surface coated by SOs under various concentrations of ATP analogs. The ATP analogs examined are presented near the data points.

### Speed and glass binding of the gliding head

We next examined the movements of gliding heads under various concentrations of ATP (Fig. 3A). The gliding speed increased with the ATP concentration, adhering simply to the Michaelis-Menten kinetics with a 1.4 μm/s maximum speed and 60 μM *K*_*m*_, the latter of which was similar to the value for ATPase activity of the jellyfish-like structure (66 μM). The gliding speed did not show an obvious sigmoidal format, suggesting that the leg does not catch SOs without nucleotides. Next, the effects of sodium azide on the gliding heads were examined (Fig. 4D). Both the number on glass and speed of gliding head decreased and reached to 4.5% and 49.6% of the original, respectively at 50 s after the addition of 154 mM sodium azide, suggesting that the activities of gliding heads were inhibited by sodium azide.

To clarify the coupling of ATP hydrolysis and displacements, we examined the effects of ATP analogs on the SO binding of the gliding head. To distinguish the SO binding involved in gliding from the non-specific binding, we examined the binding of the gliding head to solid surfaces coated by SOs, under various conditions (Fig. S4B). These results showed that the glass binding of the gliding heads occurs in a manner similar to the binding of gliding cells, suggesting that the catch of SOs by feet in different ATP forms can be monitored through the binding of gliding heads to the glass. Then, the gliding heads were mixed with various concentrations of ATP, ADP, ADP-V_i_, or ADP-AlF_x_, inserted into a tunnel of glass slides, and traced to determine the number of bound gliding heads (Fig. 4E). The numbers of bound gliding heads at 10 min after the insertion are presented as the ratio to that in 1 mM ATP. The presences of ADP-V_i_ and ADP-AlF_x_ decreased the binding activity to 35% and 50%, respectively, but ADP increased it to 220% of that without ADP, at 3 mM ADP. These results show that the leg catches SOs in the ADP form, and releases them in the ATP and ADP+P_i_ forms.

## DISCUSSION

### Gliding machinery

Although several structures involved in the gliding machinery have been visualized by conventional EM methods (22, 23, 41, 42), the positional relationship among the surface structure, the cell membrane and the jellyfish-like structure has remained unclear. The present study using ECT was able to resolve this issue, because the ECT analysis allowed three-dimensional visualization of the cells under natural conditions (Fig. 2) (43). Many flexible filaments were observed on the surface of the gliding machinery, with a length ranging from 60 to 100 nm and featured with a thick end. These are consistent with the characteristics reported for isolated Gli349 molecules, which are 97 nm long and shaped like eighth-notes in music, each with terminating globule that is designated a foot (42). It is likely that the legs observed in the present study was composed of Gli349 molecule. These molecules on the cell surface appeared flexible and did not show obvious alignment with the direction of the cell. The slice images from ECT and their average showed that the tentacle structures positioned beneath the cell membrane of the neck part at 12 nm distance and are bridged to the cell membrane through the two novel bridges (Fig. 2H arrow heads). Probably, these novel bridges hold the internal jellyfish-like structure near the membrane, and may play roles to transmit the force generated in the internal structure to the cell surface. The tentacle structures observed in negatively-stained EM images tend to appear as bundles (Fig. 3B), suggesting that they are bundled laterally by an unknown structure. The lateral protrusions found in the averaged image of tentacle shown by triangles in Fig. 2H may be responsible for this bundling. A novel periodical undercoat was found at the head part, suggesting that this undercoat connects the bell, the front part of the jellyfish-like structure to the front end of the cell membrane.

The three-dimensional image of a whole cell revealed the precise number of tentacle structures, the number of particles per tentacle structure, the length and diameter of the tentacle structures, the size of the particles, and the interval between the particles. In a previous study, the diameter and interval of the particles on the tentacle structures were reported to be 20 and 30 nm, respectively, in good agreement with the present study (22). The number of particles was estimated to be 113 to 225 per cell in the previous study (22), although we estimated it to be about 480 based on the results that the individual cells had 28 tentacle structures containing 17 particles in the present study. This difference may have been due to damage to the tentacle structures during the process of the negative staining in the previous study. The number of Gli349 molecules has been estimated as 450 per cell, based on titration using a monoclonal antibody in a previous study (19), in good agreement with the number of particles estimated in the present study. Therefore, a single leg may be driven by a single particle on the tentacle structure.

### Updating possible gliding mechanism

Previously, we proposed a working model to explain the gliding mechanism, including a repetitive cycle consisting of catch, pull, and release of SOs by the foot composed of Gli349, based on previous data as listed below (3, 5, 37). (a) On the machinery surface, large proteins, Gli349 and Gli521 work as leg and crank, respectively (41, 42). (b) Foot, the distal domain of Gli349 has binding activity to SOs (31). (c) Gliding is driven by strokes 70 nm long, 1.5 pN in force, and its direction is slightly tilted from the cell axis (7, 27, 28, 32). (d) The propelling force is used also to detach foot from SOs after stroke and foot generates drag before detach (15, 18, 34). (e) Foot can be detached to forward direction with 1.8 times less force than to backward direction (44). (f) ATP is required to detach foot from SOs, as shown by the gliding ghost experiments (29). (g) Foot catches SOs after thermal fluctuation with some cooperativity (32-34).

In the present study, the experimental data showed structural and functional changes of gliding machinery coupled to ATP hydrolysis. First, the intervals between the particles on the tentacle of jellyfish-like structure were about 5% longer in the presence of ATP related “analogs” including AMPPNP, ADP+AlF_x_, ADP+V_i_ than those in ADP and none (Fig. 3C). Second, the binding activity of gliding head, probably occurring on foot of Gli349 to SOs was accelerated by ADP than ATP-related analogs. This observation is consistent with the results of previous studies that gliding cells stayed on the glass when they were permeabilized by Triton (29). Probably, the gliding motors stayed in the ADP form under this condition. These changes in binding activity are likely linked to the structural changes found in the jellyfish-like structure. Actually, we found novel bridges linking the jellyfish-like structure to the cell membrane (Fig. 2E right, H arrow head).

Here, we update our model, based on the new information obtained in the present study (Fig. 5). (i) The unit in the ADP+P_i_ form catches SOs at a front position after thermal fluctuation. (ii) The force applied to the leg from the front triggers the release of P_i_, resulting in tight binding and force generation in the tentacle structure. The generated force transmits to the foot through several different proteins, stabilizes binding, and pulls the cell forward (stroke) to a direction slightly tilted from the cell axis. (iii) Cell movements occurring as a result of other legs pull the units forward and set back to the initial conformation for the parts other than the leg. The conformational change transmitted to the motor releases ADP. (iv) A new ATP molecule comes to the motor and allows the foot to release the SOs. The foot can be removed preferentially in the forward direction, and this directed binding property causes directed movement with a directed stroke. Then, the detached foot rebinds to SOs at the proper position after conversion from the ATP to ADP+P_i_ forms and some thermal fluctuation.

**FIG 5.**
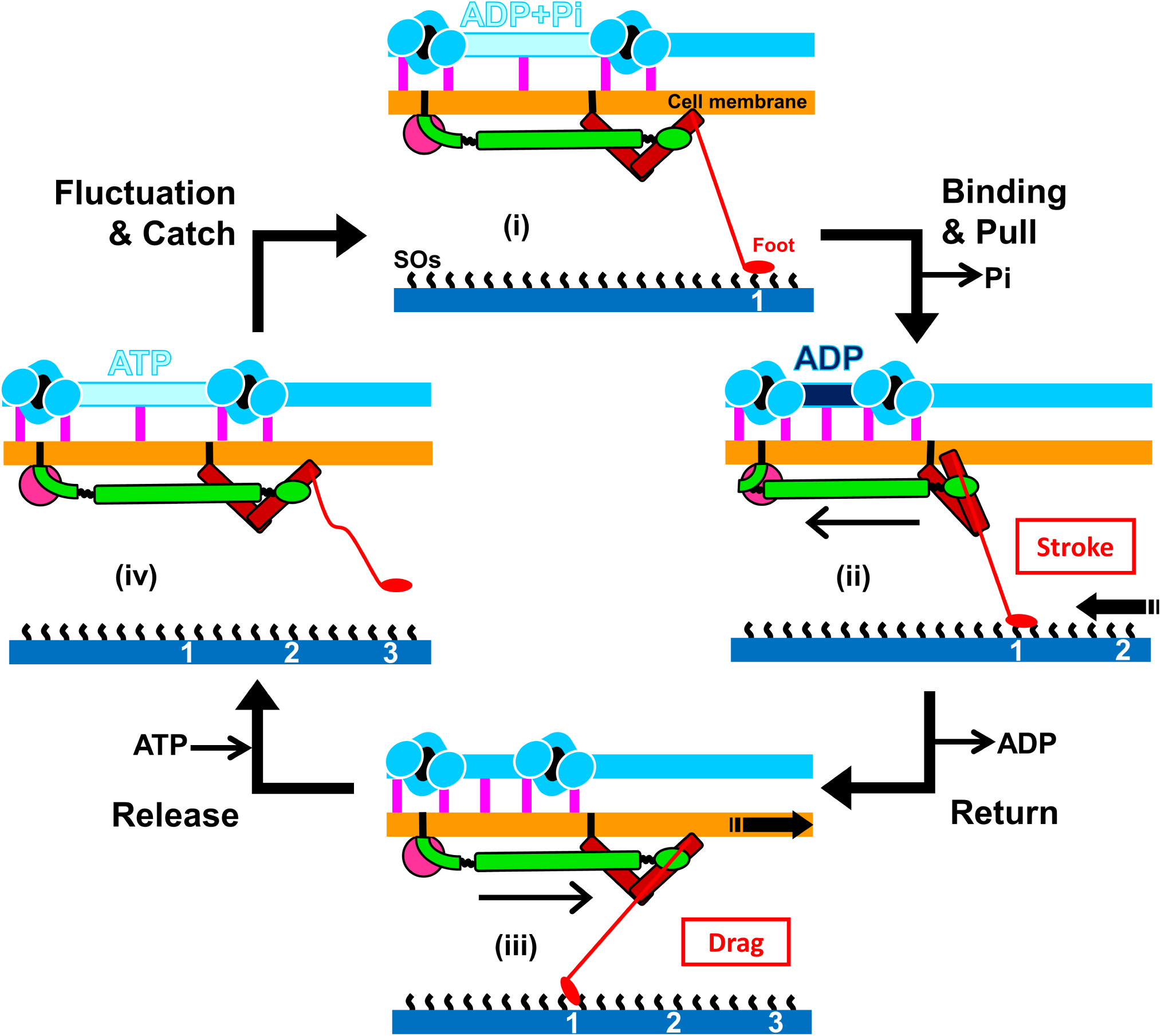
Schematic illustrations of the updated working model for the gliding mechanism. Gli123, Gli521, and Gli349, essential for gliding are presented in magenta, green, and red, respectively. The bridge between jellyfish-like structure and cell membrane is shown by pink vertical bar. The structures are not in scale to emphasize displacements. The particle interval is longer in the ATP and ADP+P_i_ forms than in the ADP forms. The foot releases and catches SOs under the conditions of ATP and ADP+P_i_ and ADP, respectively. Reactions occurring on the machinery and force exerted to the solid surface are presented by black arrows and red characters in a box. The gliding occurs through stages (i) to (iv). See the text for details.

### Perspectives

The model discussed here is consistent with all data obtained so far, but we need more information to discuss in a concrete way. First, we need images of better resolution including the jellyfish-like structure itself, as well as the structures around the cell membrane. We do not know even the assembly of three large proteins on the surface, although the individual structural outlines have been visualized (41, 42, 45, 46). Second, the conformational changes occurring with the ATP hydrolysis should be traced including cooperativity in the machinery units.

Here, we found ATPase activity in the fraction of jellyfish-like structure. This activity should be linked to the gliding motility, based on the following observations. (a) The affinities to ATP estimated from Michaelis-Menten kinetics were in similar ranges for the ATPase activity of the jellyfish-like structure and the gliding speed of the head (Fig. 3A). (b) The ATPase activity was also inhibited by sodium azide even with low affinity about hundred mM *K*_*d*_ as observed in gliding head, because azide is known to react specifically to the catalytic subunits of the F_1_-ATPase β-subunit and its related proteins featured by a P-loop or a “Walker A” motif (47). These characters were comparable to the inhibitory effects on binding and gliding speed of gliding head (Fig. 4D). (c) The jellyfish-like structure changes the conformation depending on the bound ATP analogs (Fig. 3C). (d) Among the 21 proteins identified in the gliding head, only the ATP synthase paralogs of the jellyfish-like structure can be suggested for ATPase from amino acid sequence (Fig. S4A(i)).

If the ATPase activity is derived from the β subunit paralog, its maximum activity can be calculated as 0.09 turnover /s. This number can provide the gliding speed of 4.5 μm per s, if we assume that mycoplasma cells can glide with a stroke of 70 nm coupled with the ATP hydrolysis, occurring in one of the 450 legs (19). Alternatively, the ATPase activity may be stimulated when the legs catch SOs in the gliding mechanism. If the catalytic subunits of F_1_-ATPase paralog play the central roles in the gliding mechanism, this fact may suggest that the gliding mechanism was evolutionally originated from the combination between F_1_-ATPase and adhesin. It also suggests a novel evolutional destination for F_1_-ATPase. (3, 5, 25).

## MATERIALS AND METHODS

### Strains and culture conditions

A mutant strain of *M. mobile*, P476R *gli521*, which binds to SOs more tightly than the wild-type strain, was used in this study (7, 14, 18, 29). Cells were cultured as previously described (21, 48). The non-gliding mutant strains, m13 and m23 mutated for the *gli349* gene were reported previously (7, 15, 18, 21).

### Optical microscopy

Mycoplasma cells and cell fractions were observed by phase-contrast microscopy using an IX71 microscope (Olympus, Tokyo, Japan). The image and movement were recorded with a Wat-120N charge-coupled-device (CCD) camera (Watec, Yamagata, Japan). All video data were analyzed by using ImageJ software, version 1.7 (http://rsb.info.nih.gov/ij/). Fluorescence microscopy was performed for the signals by EYFP fusion and immunofluorescence, as previously described (8, 24).

### EM

For ECT of the permeabilized cells and the jellyfish-like structure, cultured cells were collected by centrifugation at 12,000 × *g* for 4 min at RT and suspended in a fresh growth medium at a 1500-fold-higher concentration. A cell suspension of 1.5 μl was placed on an EM grid for 10 min, and 1.5 μl detergent solution (1.4% Tween 20 or 0.16% Triton X-100, 1 mg/ml DNase, and 5 mM MgCl_2_ in phosphate-buffered saline (PBS) consisting of 75 mM sodium phosphate (pH 7.3) and 68 mM NaCl) was added. In the case of the gliding heads and intact cells, 2.6 μl suspensions derived from 150 and 15 ml cultures, respectively, were applied to the EM grids. The images were observed and captured at 0.570 nm per pixel by a Titan Krios FEG transmission electron microscope (FEI, Eindhoven, Netherlands) operated at 300 kV on an FEI Falcon 4 k × 4 k direct electron detector (FEI). Single-axis tilt series were collected covering an angular range from -70° to +70° with a nonlinear Saxton tilt scheme at 5–10 μm underfocus using the Xplore 3D software package (FEI). The other experimental procedures were performed as previously described (49, 50). Images were generally binned two-fold and 3D reconstructions were calculated using the IMOD software package (51). Fourier transform was calculated by ImageJ 1.37v (http://rsb.info.nih.gov/ij/). For negatively-stained EM, the jellyfish-like structures were prepared on the grid as previously described (22). To examine the effect of ATP analogs, the Triton solution was removed after washing by buffer A (2 mM MgCl_2,_ 20 mM Tris-HCl at pH 7.5), and the jellyfish-like structure was treated with buffer A containing an ATP analog at 2 mM, ADP, AMPPNP, ADP-V_i_, or ADP-AlF_x_. The grid was stained for 1 min by 2% ammonium molybdate (wt/vol) and air-dried (22, 52). The specimens were observed by a transmission electron microscope, JEM-1010 (JEOL, Tokyo, Japan) at 80 kV, equipped by a FastScan-F214 (T) CCD camera (TVIPS, Gauting, Germany). Image averaging was done by EMAN, version 1.9 (http://ncmi.bcm.tmc.edu/_stevel/EMAN/) for both EM methods. Surface-rending images were obtained using the three-dimensional modeling software Amira 5.2.2 (Visage Imaging, San Diego, CA).

To analyze the change in the periodicity of the chain structure, the negatively-stained images were captured at 0.744 nm per pixel (Fig. S2). More than hundred images 45 pixel wide and 216 pixel long including a tentacle were manually selected for each ATP analog. The obtained images were classified and averaged into nine classes. The averaged images were profiled for a rectangle 18 pixel wide and 83 pixel long. The distance between two negative peaks corresponding to the center of particles were measured for all classes and averaged.

### ATPase assay

Mycoplasma cells were collected by centrifugation at 12,000 × *g* for 10 min at RT, washed twice with PBS/G (PBS containing 10 mM glucose), and suspended in PBS/G to be 50-fold concentrated from the culture. The cells were treated with Triton solution (0.3% Triton X-100, 1 mg/ml DNase, and 5 mM MgCl_2_ in PBS) for 1 min at RT. The Triton-insoluble fraction was collected by centrifugation at 20,000 × *g* for 20 min at 4°C, and suspended in buffer A. This process was repeated. ATPase was assayed by a continuous spectrophotometric method using a 2-amino-6-mercapto-7-methylpurine ribonucleoside/purine nucleoside phosphorylase reaction to detect the released inorganic phosphate (EnzChek Kit; Life Technologies, California, USA) (53). The reaction conditions were as follows: 20 μg/ml jellyfish-like structure, 2 mM MgCl_2_, 20 mM Tris-HCl (pH 7.5) in a total volume of 0.2 ml, at RT. Sodium azide was added at a concentration of 15.4 or 154 mM when stated. The protein amount of the ATPase β-subunit paralog was estimated from the densitometry of SDS-PAGE.

### Gliding head

*Mycoplasma* cells were collected by centrifugation, washed with PBS/G, and suspended in PBS/G containing 0.1% Tween 60 to be 15-fold concentrated from the culture. After 30 min, the cells were applied to a sonicator (UR-20P; TOMY SEIKO, Tokyo, Japan) for 1 s, 60 times on ice. The sonicated fractions were treated with 1 mg/ml DNase, 1 mg/ml RNase and 10 mM MgCl_2_, and subjected to a stepwise gradient centrifugation consisting of 0%, 20%, 30%, 40%, 50% and 60% sucrose layers. After centrifugation at 18,000 × *g* for 15 min at 4°C, the fraction of gliding heads was recovered from the top of the 30% sucrose layer and washed with buffer A. The gliding head suspension was inserted into a tunnel at time 0, and the number of bound heads in a 32 × 43 μm^2^ field were traced under the microscope. The tunnel was assembled using two pieces of glass and precoated with 10 mg/ml fetuin in buffer A for 10 s, then washed with 50 μl buffer A (8). The numbers of heads in the three fields were averaged for each time point and presented as a ratio to the number of gliding heads in the presence of 1 mM ATP at 10 min, with standard deviation. The cell behaviors were analyzed as previously reported (8, 34). SDS-PAGE, PMF, and Western blotting were performed as previously described (22). The monoclonal antibodies used were also described previously (16, 20, 54).

## FUNDING INFORMATION

This work was supported by a Grant-in-Aid for Scientific Research on the Innovative Area Harmonized Supramolecular Motility Machinery and Its Diversity (MEXT KAKENHI Grant Number 24117002), by a Grants-in-Aid for Scientific Research (B) and (A) (MEXT KAKENHI Grant Numbers 24390107, 17H01544), by JST CREST Grant Number JPMJCR19S5, Japan, by the Osaka City University (OCU) Strategic Research Grant 2018 for top priority researches and by a Grant-in-aid of the FUGAKU TRUST FOR MEDICINAL RESEARCH to MM, and by a Grant-in-Aid for Specially promoted Research (JSPS KAKENHI Grant Number 25000013) to KN.

## ACKNOWLEDGMENTS

We appreciate the helpful input of John Heuser at Kyoto University, and Eisaku Katayama and Isil Tulum at Osaka City University.

## (FIGURE LEGENDS)

**FIG. S1. Images and analyses by ECT.** (A) Filamentous structure on the surface of a permeabilized cell observed by ECT. Original and colored images are presented in the left and right panels, respectively. The thick end of a filamentous structure is marked by a white triangle. (B) Distributions of parameters in tentacle structures analyzed by ECT. The distance of tentacle structures from the cell membrane, number, and length are shown in (i), (ii), and (iii), respectively. The average values are shown as filled triangles.

**FIG. S2. Procedure to measure particle intervals in presence of different ATP analogs.** (A) Averaging images of tentacles under individual ATP analogs. Tentacle images of straight part were manually picked, classified, and averaged. (B) Measurement of particle intervals based on density profile. See Materials and Methods for details.)

**FIG. S3. Optical microscopy of gliding heads.** (A) Occasional detachment of the gliding head from the cell body. The movie images of cells in 0.1% Tween 60 are presented for 5.67 s. The cell was gliding in the direction indicated by the blue arrow. The second gliding machinery probably in preparation for cell division began to glide in the opposite direction indicated by the red arrow. Suddenly, the second gliding machinery detached from the other part of the cell, as indicated by the yellow triangle, and glided away. A trace for 1.34 s is shown in the lower rightmost panel. This phenomenon was observed consistently with a frequency of about one per 300 cells over the 2-min observation period after the addition of Tween 60. (B) Localization of gliding and surface proteins in original cells and gliding heads examined by immunofluorescence microscopy. Fluorescent signals labeled for the proteins are overlaid onto the phase-contrast images.

**FIG. S4. Gliding head characterization.** (A) Protein components of the gliding head. (i) Protein profiles of the original cell, elongated cell, and gliding head. The proteins of each fraction shown in the right lower box were developed by SDS-10%PAGE and identified by PMF as listed in the table. The proteins specific for the gliding head are listed from a to h. Two proteins were identified from band g. Gli349 has a faster migration speed than that expected from the marker proteins. The proteins abundant in the original and the elongated cells are listed from i to t. JSP stands for jellyfish structure protein. (ii) Western blotting analyses for gliding and surface proteins. The amount derived from the same number of cells was applied to each lane to compare the protein amounts included in each fraction. (B) Time course of the binding of gliding heads to the glass surface coated by fetuin under various conditions. Non-binding mutants, m13 and m23, sialyllactose (SL), anti-Gli349 antibody (Ab), and bovine serum albumin (BSA) were examined. The number of bound gliding heads increased with time in a manner comparable to that in the original cells. The binding of gliding heads was not observed in the presence of 5 mM free SL or to the glass precoated with 20 mg/ml BSA instead of fetuin, demonstrating that the binding observed here was mediated by SOs fixed on the glass. The binding was blocked by a monoclonal antibody against Gli349 at 0.05 mg/ml, and the binding did not occur when the cell pieces were prepared from m13 and m23 mutants deficient in glass binding due to mutations in Gli349.

**Movie S1.** Gliding motility of *M. mobile* (P476R *gli521* mutant) observed by phase-contrast microscopy at real time. The horizontal edge of the movie is 80 μm.

**Movie S2.** Slice movie of an intact cell reconstructed from ECT. The horizontal edge of the movie is 793 nm.

**Movie S3.** Slice movie of a permeabilized cell reconstructed from ECT. The horizontal edge of the movie is 917 nm.

**Movie S4.** Slice movie of the jellyfish structure reconstructed from ECT. The horizontal edge of the movie is 848 nm.

**Movie S5.** Three dimensional image rendered for 146 nm-thick slice of permeabilized cell reconstructed from ECT. The horizontal edge of the movie is 848 nm.

**Movie S6.** Gliding heads. The horizontal edge of the movie is 64 μm.

